# The cost of behavioral flexibility: reversal learning driven by a spiking neural network

**DOI:** 10.1101/2024.05.16.594474

**Authors:** Behnam Ghazinouri, Sen Cheng

**Author notes:** { }. http://www.rub.de/cns.

## Abstract

To survive in a changing world, animals often need to suppress an obsolete behavior and acquire a new one. This process is known as reversal learning (RL). The neural mechanisms underlying RL in spatial navigation have received limited attention and it remains unclear what neural mechanisms maintain behavioral flexibility. We extended an existing closed-loop simulator of spatial navigation and learning, based on spiking neural networks [8]. The activity of place cells and boundary cells were fed as inputs to action selection neurons, which drove the movement of the agent. When the agent reached the goal, behavior was reinforced with spike-timing-dependent plasticity (STDP) coupled with an eligibility trace which marks synaptic connections for future reward-based updates. The modeled RL task had an ABA design, where the goal was switched between two locations A and B every 10 trials. Agents using symmetric STDP excel initially on finding target A, but fail to find target B after the goal switch, persevering on target A. Using asymmetric STDP, using many small place fields, and injecting short noise pulses to action selection neurons were effective in driving spatial exploration in the absence of rewards, which ultimately led to finding target B. However, this flexibility came at the price of slower learning and lower performance. Our work shows three examples of neural mechanisms that achieve flexibility at the behavioral level, each with different characteristic costs.

## 1 Introduction

Spatial navigation and reversal learning (RL) are fundamental skills for mobile animals. Spatial navigation involves identifying and maintaining a path between two locations [15]. Research over the past 50 years has identified key brain regions for spatial navigation in the medial temporal lobe and spatially selective cells like place cell (PC) [14], boundary cells (BC) [10], and head direction cells [17]. Recent studies also point to the hippocampus’ role in RL during spatial navigation tasks [19], where an obsolete behavior has to be suppressed and a new one has to be acquired. Recent results suggest that replay might be involved in the neural mechanism underlying RL [6]. While many computational models have been developed for spatial navigation, they focus on static environments [3, 4, 8, 12] and the dynamic aspects of RL have received limited attention.

We focus on an ABA RL task where an agent has to learn to navigate to a fixed target (A), then to a different target (B), and finally to target A again. This task touches on two trade-offs: stability vs. plasticity, well known in the neural network literature [18], and exploitation vs. exploration, which is central in the reinforcement learning literature [16]. For the RL task, maintaining behavioral flexibility requires exploration and plasticity, but this reduces performance and stability. The challenge is understanding how a biologically plausible spiking neural network maintains flexibility and its associated costs.

Using a spiking neural network, we find that a combination of symmetric spike-timing dependent plasticity (STDP) and optimized place field parameters performs well on the first target, but lacks flexibility for the second. In three other cases, the agent remains flexible, but incurs different costs. Asymmetric STDP results in highly variable behavior. Using many small place fields leads to low overall performance. Providing an external supervisory signal (injecting noise when unrewarded for too long) results in slow RL and variable performance on the second target, but better performance on the first. Our results suggest multiple neural mechanisms can lead to behavioral flexibility, each with a distinct profile, allowing biological agent to evolve or develop solutions that fit its needs.

## 2 Methods

To model RL in a biologically plausible network model, we adopted the CoBeL-spike (Closed-Loop Simulator of **Co**mplex **Be**havior and **L**earning Based on Spiking Neural Networks) framework^1^ [8].

### Behavioral task

The spatial navigation task was similar to the Morris water maze [13]. Simulations were conducted in a 2.4 m *×* 2.4 m open field. Each simulation included 30 trials, in which the agent started at the center [*x*_0_, *y*_0_, *t*_0_] = [1.2, 1.2, 0.0] and freely navigated to find a hidden target, a circular area (diameter= 0.4 m). A trial ended when the agent reached the target or after 5 seconds. Upon reaching the target, the agent received a reward, aiding learning for quicker subsequent trials. In the first phase (trials 1-10), the target was at [0.5, 0.5] (target A). The agent learned to find this target from the start location. In the second phase (trials 11-20), the target moved to [−0.5, −0.5] (target B), requiring the agent to unlearn the previous location and learn the new one. In the third phase (trials 21-30), the target returned to location A, and the agent had to either remember the original location or re-learn it.

### Overview of the computational model

The CoBeL-spike toolchain comprises two components: a virtual environment and a learning agent (Fig. 1). The virtual environment, an open field Gym environment powered by OpenAI [2], monitors the agent’s location and ensures that actions adhere to physical limitations. At each 0.01 s time step, the environment transmits the current trial time, Cartesian coordinates, and reward signal ([*x*_*i*_, *y*_*i*_, *t*_*i*_],[*r*_*i*_]) to the agent via ZMQ (**Z**ero **M**essage **Q**ueuing) [11] to MUSIC (**MU**lti **SI**mulator **C**oordinator) [5]. MUSIC integrates the Gym environment with the learning agent, enabling real-time data transmission.

**Fig. 1.**
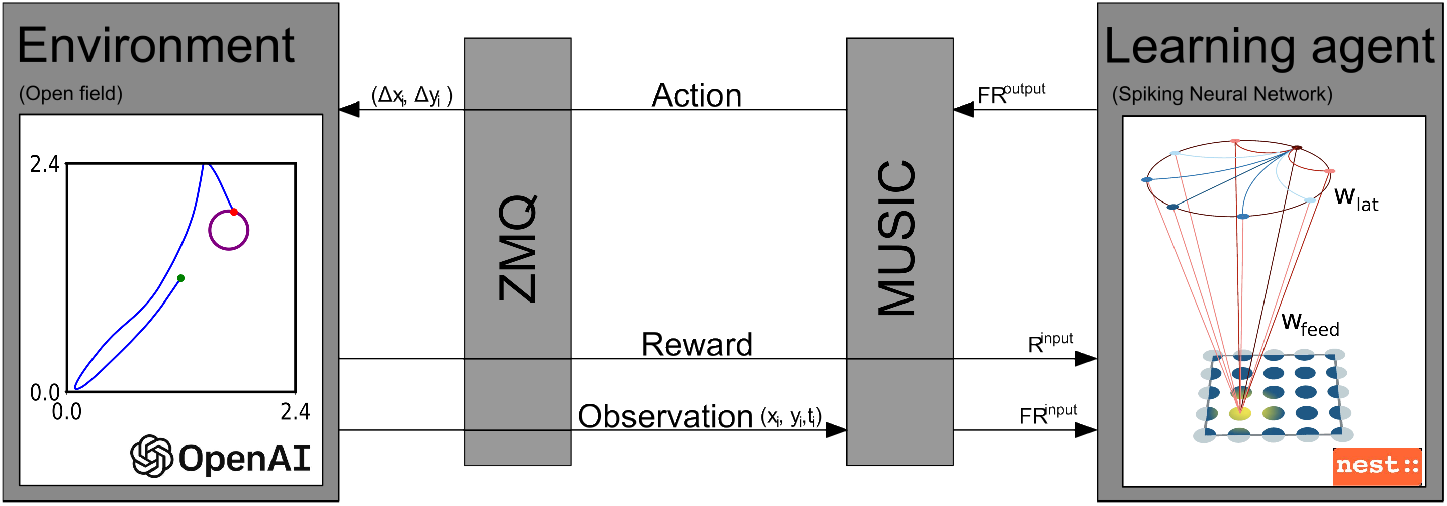
Toolchain components. The CoBeL-spike toolchain consists of two components: a virtual environment (an OpenAI Gym environment) and a learning agent (two-layered spiking neural network). ZMQ and MUSIC provide interfaces between the two model components. For details see the main text.

The learning agent is a two-layer spiking neural network implemented in NEST (**NE**ural **S**imulation **T**ool) [7]. The network’s first layer comprises PC, BC, and nose cells (NC). At each time step, MUSIC calculates the firing rates of these cells: (*x*_*i*_, *y*_*i*_, *t*_*i*_) −→ Ω_*PC*_, Ω_*BC*_, Ω_*NC*_. The output layer contains 40 action selection neurons, whose activity is mapped to movement in Cartesian space ([Δ*x*_*i*_, Δ*y*_*i*_]) by MUSIC via ZMQ to the environment. The action selection neurons drive movement and may represent the striatum [9]. The environment updates the agent’s location based on the action, provided the new location is valid; otherwise, the agent remains in place. In the example trial shown in Fig. 1, the agent started at the center (green marker), moved through the environment (blue curve), and reached the target (purple circle).

### Simulation of neural activity

Each unit in the first layer consists of a Poisson generator and a parrot neuron [7]. The number of PC (*N*_*PC*_) varies across simulations. The firing fields of PC are evenly distributed across the 2D environment. Each firing field 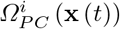 is a Gaussian 2D distribution with width *σ*_*PC*_ [8]. Each PC has an excitatory connection to all actions selection neurons in the second layer. These feedforward weights are plastic (see below, with initial weights *W* ∼ *N* (30, 5).

To prevent the agent from getting stuck at the edges and corners of the environment, the first layer has eight BC [10] that project to the second layer.

These cells drive the agent towards the environment’s interior. Each BC is active 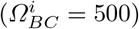 when the agent is inside its field [8].

In some simulations, we examined the effect of noise on the agent’s performance. In this case, noise cells (*N*_*NC*_ = 40) were added to the network with static one-to-one excitatory connections to the second-layer neurons. Noise is introduced by activating a random set of five neighboring noise cells at time steps 2.5 s, 3.5 s, and 4.5 s, with each activation lasting 0.4 s.

The second layer consists of 40 action selection neurons modeled as Leaky Integrate-and-Fire (LIF) [7]. Each neuron represents a movement direction evenly distributed across 360^*°*^. These neurons form a continuous ring attractor network [3, 8], with local excitation and global inhibition [8].

### Synaptic plasticity and learning

The synaptic weights between the first and second layers significantly impact the agent’s behavior and performance. Only the weights between place cells and action selection neurons are plastic. We applied STDP with an eligibility trace (*stdp_dopamine_synapse* in NEST [7]).

The potential weight changes depend on the temporal difference between a post- and a pre-synaptic spike, Δ*t* = *t*_*post*_ − *t*_*pre*_,

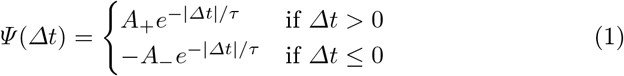

where *A*_+_ and *A*_−_ are the amplitudes of the weight change, and *τ* is a time constant. In all simulations *A*_+_ = 0.1 and *A*_−_ = 0.1 for asymmetric STDP and *A*_−_ = −0.1 for symmetric STDP. The symmetric STDP rule results in long-term potentiation (LTP) regardless of spiking order, whereas the asymmetric STDP leads to LTP only, if the post-synaptic spike follows the pre-synaptic spike, otherwise it leads to long-term depression (LTD).

In addition to the relative time difference between pre- and post-synaptic spikes, the weight change is also a function of the eligibility trace, which increases, if the agent finds the target location and dopamine is released and then decays exponentially [7]. The latter is critical in ensuring that later place cells (those activated closer to reaching the target) have a larger synaptic update.The reward signal is sent to the dopaminergic neurons through MUSIC.

### Measuring the agent’s performance

To evaluate the agent’s navigation performance, we measured three aspects of its spatial trajectory: escape latency, proximity to the target, and similarity to the ideal trajectory. Escape latency is the time from the start of a trial to when the agent finds the target, with a maximum of 5 s for timeout. This was our default measure due to its simplicity and common use in animal experiments. Additionally, we captured cases of partial learning (the agent approaches, but does not reach, the target) by measuring the minimum distance between the agent’s trajectory and the target.

However, there are differences between the trajectory of the agent and the ideal path that neither latency nor minimum distance can capture. We therefore quantified the similarity between the agent’s trajectory and the ideal path by using dynamic time warping (DTW), a technique for measuring the similarity between two sequences, which may differ in length, speed or timing [1]. Given two sequences *X* = {*x*_1_, *x*_2_, …, *x*_*n*_} and *Y* = {*y*_1_, *y*_2_, …, *y*_*m*_}, DTW constructs an *n × m* distance matrix *D*, where each element *D*[*i, j*] represents the distance between *x*_*i*_ and *y*_*j*_, typically calculated using the Euclidean norm. The accumulated cost matrix *C* is computed as follows:

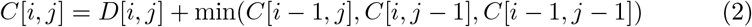

This recurrence relation determines the minimal cumulative distance to align the subsequences *X*[1 : *i*] and *Y* [1 : *j*]. The optimal warping path is the monotonic path from [1, 1] to [*n, m*], which always moves from one element of *C*[*i, j*] to the neighboring one with the lowest value. Figure 2 demonstrates DTW between a sample trajectory and two optimal paths to targets A and B. The grey lines connect the points along the optimal warping path. The DTW distance between the two sequences is given by *C*[*n, m*].

**Fig. 2.**
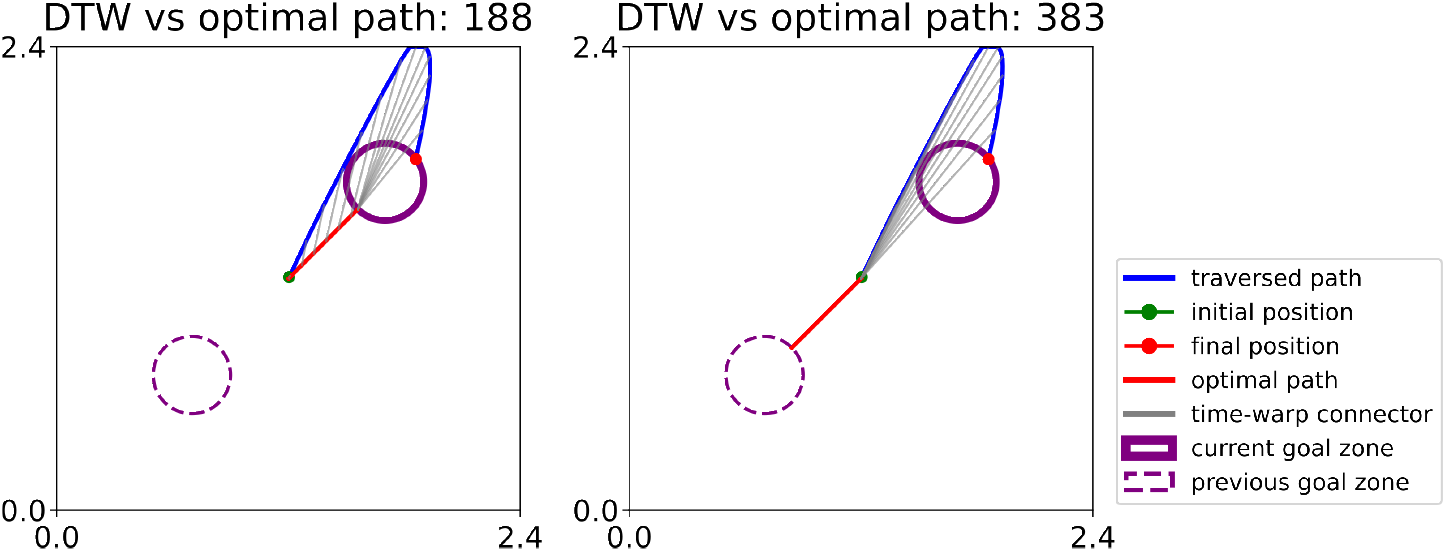
Illustration of dynamic time warping (DTW). DTW quantifies the similarity between the agent’s trajectory in a trial (blue curve) and the optimal path (red line). To study which target the agent is most likely to aim for, we compare DTW for navigation towards the current target (left) and the previous target (right). Gray lines represent the time-warp relationship. A path closer to the ideal has lower DTW value.

## 3 Results

### 3.1 Studying behavioral flexibility in a learning agent based on spiking neural networks

To investigate behavioral flexibility in RL in spatial navigation tasks, we utilized the CoBeL-spike toolchain, focusing on the role of the STDP rule and the properties of PC, such as field size (*σ*_*PC*_) and number of PC (*N*_*PC*_). Additionally, we examined how this process can be manipulated by external factors.

As the baseline agent, we chose parameters (symmetric STDP, *σ*_*P C*_ = 0.2 *m* and *N*_*P C*_ = 21^2^) that were most effective in the task with a single target [8]). The agent successfully learned to navigate to target A efficiently after 10 trials (Fig. 3A). However, in the second phase of the ABA RL task, the agent consistently returned to target A and failed to find target B. Performance improved again once the goal shifted back to target A. Most features of spatial learning are captured by the learning curves based on the average escape latency (Fig. 4A, top) The agent learned to navigate to target A within the first 10 trials, but the escape latency in the second phase increased to nearly 5 s, the time-out value. Good performance returned in the third phase, when the goal was target A again. To confirm the observation from the single examples that in the second phase the agent persevered on target A, we analyzed the trajectories of the agent in the second phase (Fig. 4A, middle and bottom). Proximity and DTW metrics with respect to the current target B are represented by a solid green line, and to the former target A by a dashed green line. Indeed, the agent continues to seek target A in the second phase, even though this is not rewarded.

**Fig. 3.**
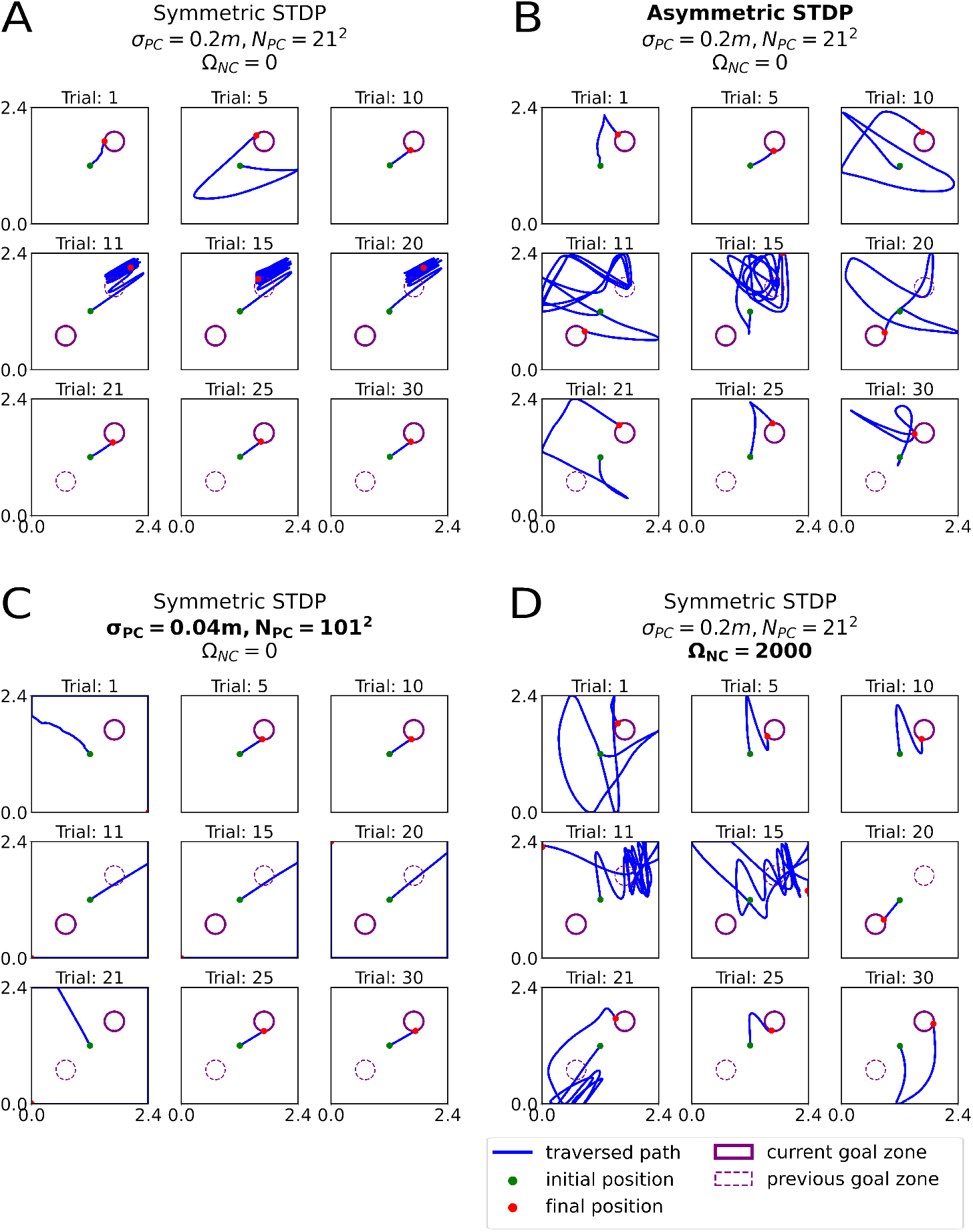
Behavior in simulated RL task. A-D: Example trajectories from four simulations, each comprising 30 trials split into three phases (rows). Within each row, plots illustrate trials 1, 5, and 10. In each trial, the agent starts at the center of the environment, seeks and learns the target (solid purple circle). The target is located in the top-right corner (target A) in the first and third phases and in the bottom-left corner (target B) in the second phase. A dashed circle marks a former target location. **A:** Symmetric STDP. **B:** Asymmetric STDP. **C:** Symmetric STDP with many smaller fields. **D:** Symmetric STDP with noise pulses injected to the action neurons.

**Fig. 4.**
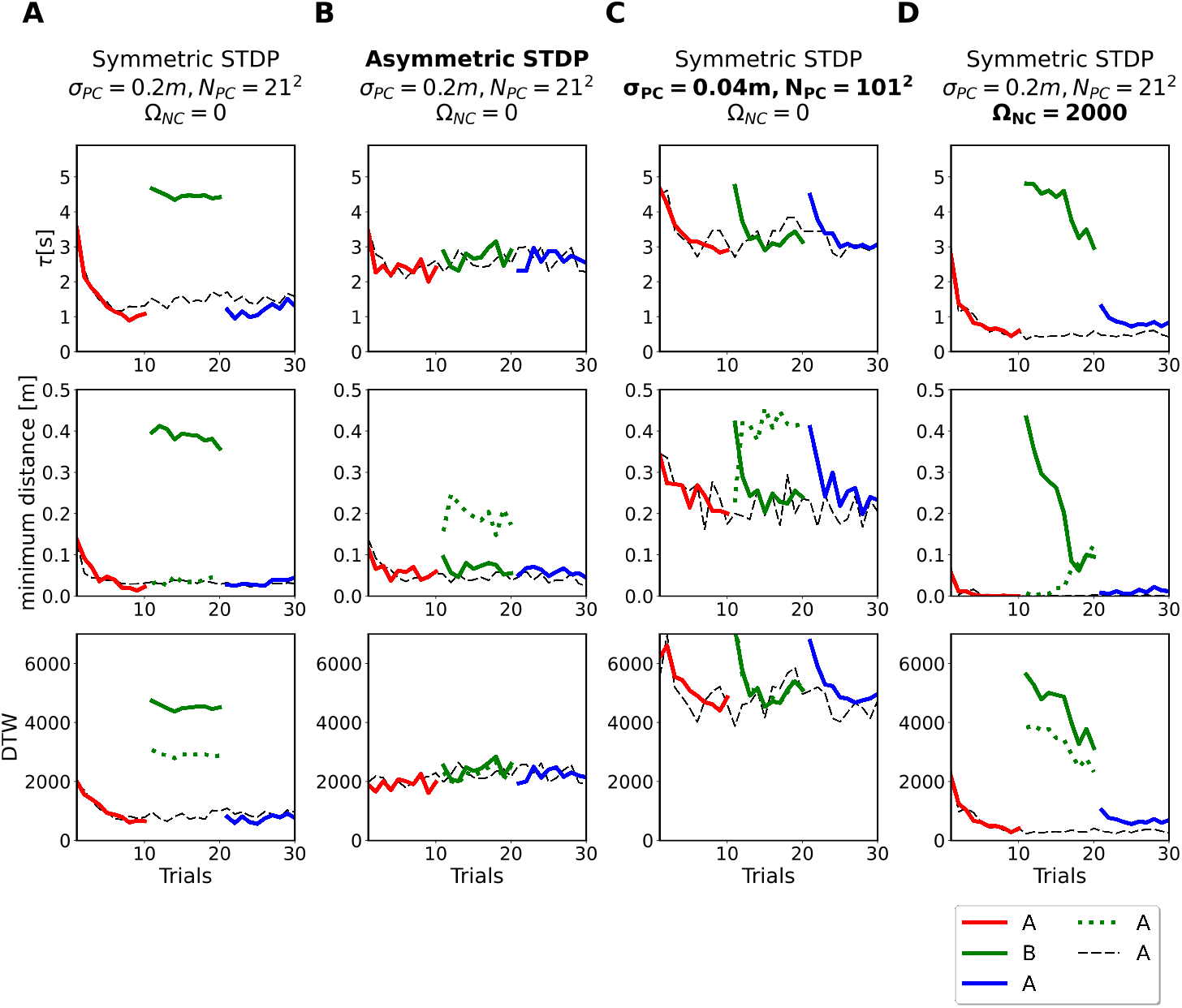
Learning curves during RL. A-D: Each panel illustrates different measures of learning for three phases of an RL task (represented by red, green, and blue curves, respectively) compared to a baseline task with no reversal (dashed black line). Results were averaged over 50 repetitions. Top: Escape latency. Middle: Dynamic-Time-Warping (DTW). Additionally, DTW measure is presented for target A (dashed green line) in the second phase, when the goal was target B (solid green line). Bottom: Minimum distance to the current target (solid lines) or former target (dashed green line). **A:** Symmetric STDP. The agent learns target A quickly during the first phase, but fails to explore and learn target B after the target switch. **B:** Asymmetric STDP. The agent learns the location of the target A in the first phase and target B in the second. **C:** Symmetric STDP with many smaller fields. In each phase, the agent adapts to a new target and unlearns the previous one. **D:** Symmetric STDP combined with noise injections. The agent learns target B in the second phase.

We hypothesized that the baseline agent lacks behavioral flexibility for two main reasons. 1. The behavior of the baseline agent is focused on exploitation and neglects exploration. 2. Symmetric STDP can only potentiate synapses and a new behavior could only emerge, if the new association was stronger the previous ones. However, the established associations prevent new behaviors to emerge (stability).

### 3.2 Network features that introduce behavioral flexibility

To test our hypotheses, we introduced an asymmetric STDP learning rule while the other parameters remained identical to the baseline agent. This agent is indeed more flexible, learning target B (Fig. 3B) and performing much better than the baseline agent in the second phase (Fig. 4B). However, this flexibility comes at a cost — this agent’s performance is reduced during the first phase compared to the baseline agent (Fig. 4A,B).

We confirmed this pattern across varying place cell parameters for symmetric (Fig. 5A) and asymmetric (Fig. 5B) STDP rules. We scaled the field size, the PC number, or both simultaneously. In all scenarios, agents with symmetric STDP performed better on target A, while those with asymmetric STDP reached target B mostly independent of place cell parameters. The lower performance is the result of more variable behavior, as is evident in the example trajectory (Fig. 3B), the higher average latency and higher DTW, but the minimum distance is not worse (Fig. 4B). Hence, the behavioral flexibility afforded by the asymmetric STDP comes at the cost of more variable behavior.

**Fig. 5.**
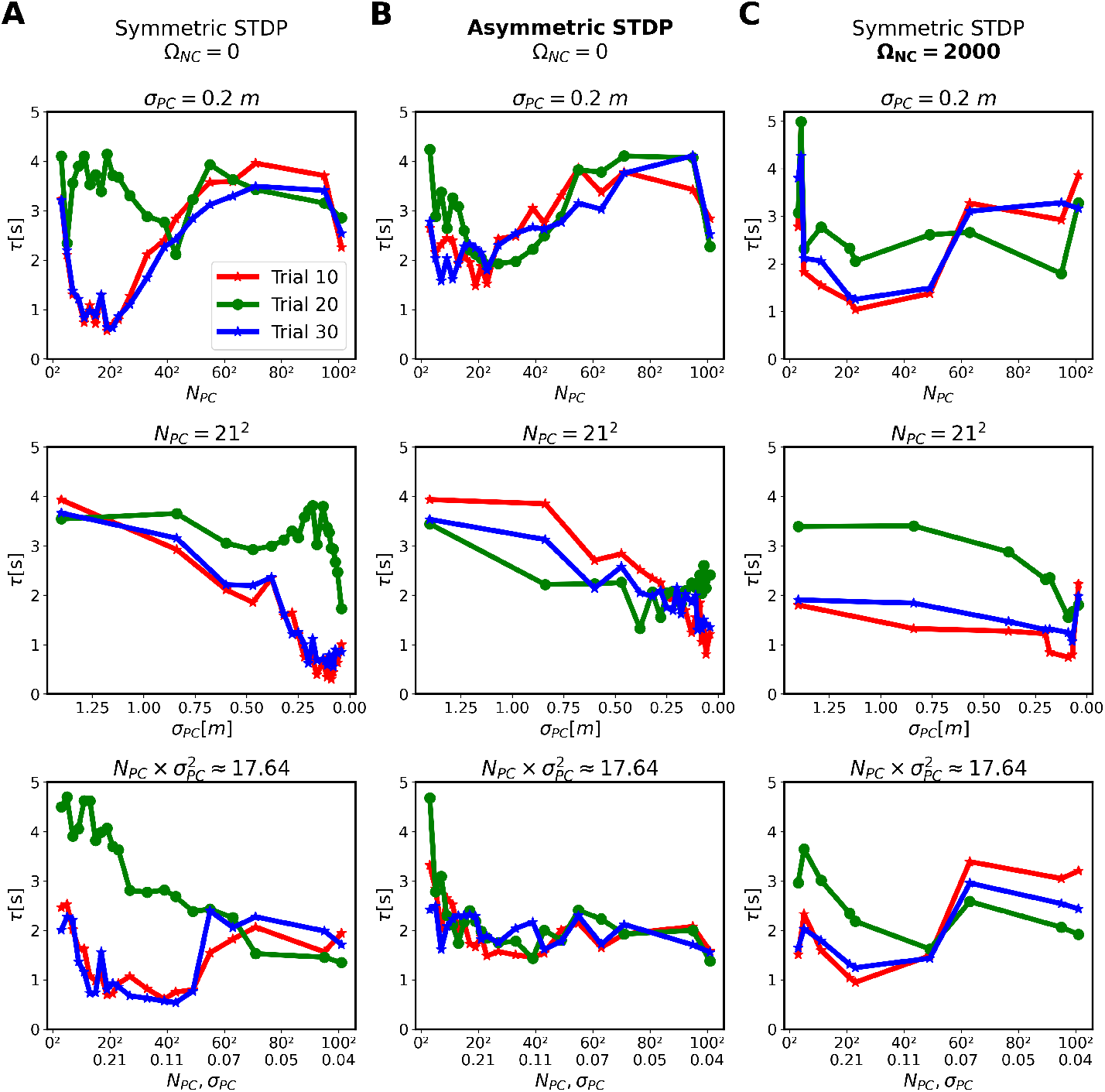
Dependence of RL on plasticity rule, place cell parameters and external noise. Each panel shows the average escape latency across 30 simulations on the last trial in each phase (red: first phase, green: second phase, blue: third phase) as a function of some place cell parameters for **A**: symmetric STDP, **B**: asymmetric STDP, and **C**: symmetric STDP with noise. Each row represents different scaling of place cell parameters. Top: Scaling place cell number *N*_*PC*_, while keeping field size fixed *σ*_*PC*_ = 0.2 m. Middle: Scaling *σ*_*PC*_, while keeping *N*_*PC*_ fixed. Bottom: Simultaneous scaling of both variables, such that 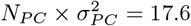.

Close inspection reveals that using symmetric STDP, the agent did perform well on target B, if there are many and small place fields (Fig. 5A, bottom). We confirmed that, in the case *N*_*P C*_ = 101^2^ and *σ*_*P C*_ = 0.04 *m*, the agent with symmetric STDP indeed explored and learned target B (Fig. 3C). However, the agent performed more poorly at target A compared to the baseline agent on all three learning measures (Fig. 4C). Even more dramatically, the agent almost forgot target A in the second phase and had to relearn it in the third phase as if it was naïve. This indicates that the behavioral flexibility, in this case, comes at the cost of destabilizing the already learned information.

### 3.3 Driving behavioral flexibility through an external signal

Another alternative is using an external supervisor to detect when the agent has not been rewarded for too long and inject noise into the network (Ω_*NC*_ = 2000, see Methods) to disrupt repetitive loops and promote exploration (Fig. 3D, trials 11 and 15). In the third phase, the agent initially gets stuck at target B, but escapes this loop to find target A again. Comparing learning curves with and without noise injection shows that noise significantly enhanced performance in the second phase (Fig. 4A and D). However, this improvement varied across place field parameters (Fig. 5C).

Systematic analysis of different noise magnitudes indicated that performance improved with increased noise firing rate up to a point (Fig. 6). Beyond this, additional noise did not further boost performance. Notably, noise injection also improved performance in the first and third phases (Figs. 4D and 6), suggesting that noise helps the agent explore more thoroughly and find more efficient trajectories, similar to the benefit of the *ε*-greedy strategy in reinforcement learning.

**Fig. 6.**
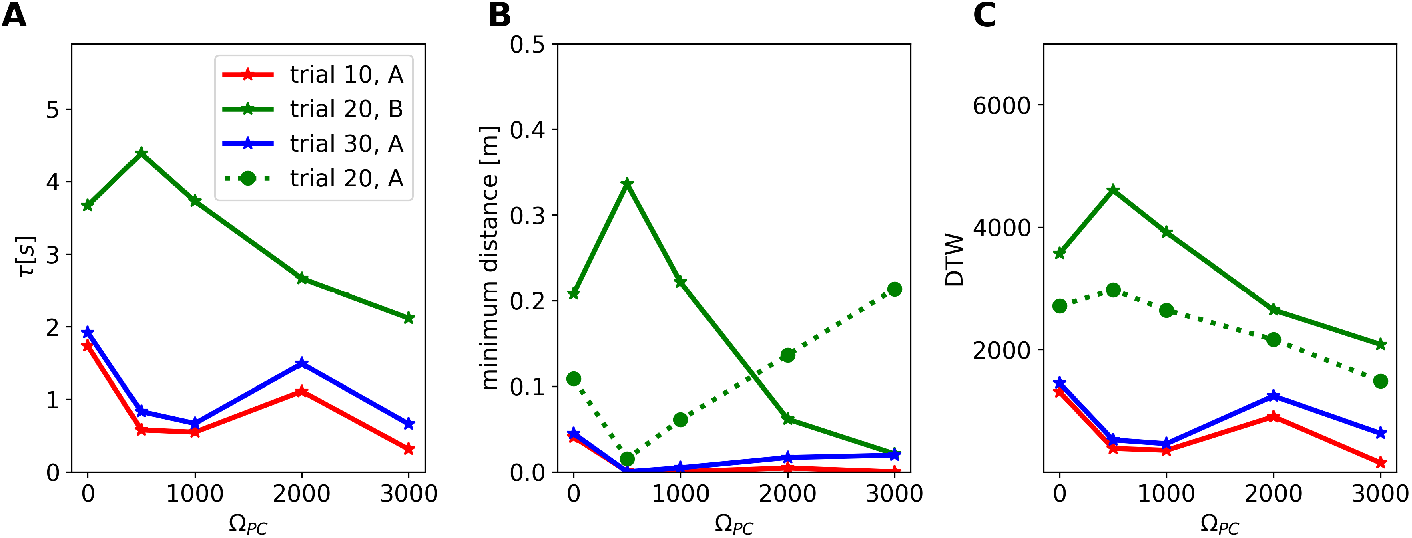
Effects of noise injection during ABA RL. **A**: Average escape latency, **B**: the minimum distance to the target, and **C**: DTW on the last trial of each phase (trials 10, 20, and 30) as a function of the firing rate of the noise cells (Ω_*NC*_). In each trial, noise impulses were injected at 2.5*s*, 3.5*s*, and 4.5*s*. Each impulse had a duration of 0.4*s* and was randomly injected into a cluster of five adjacent AC neurons.

## 4 Discussion

We modeled reversal learning (RL) in spatial learning using spiking neural networks in a closed-loop simulation with reward-driven STDP. An agent with symmetric STDP and optimized place cell properties excels at learning a specific target, but struggles to unlearn it, explore, and learn new targets. The agent’s behavior is dominated by exploitation and stability at the expense of exploration and plasticity. To encourage exploration and reduce behavioral stability, we found three solutions, each with its own costs: applying asymmetric STDP, using many small place fields, and injecting external noise.

The closest simulation frameworks to ours are those our model is based on [3]. While that paper touched on goal relocation, it primarily focused on sequential neuromodulation of synaptic plasticity rules in learning navigation tasks, not on RL. Another framework simulates closed-loop behavior using reinforcement learning but emphasizes the role of context in ABA extinction learning [20] and is less biologically plausible compared to ours.

There is an analogy to attractor dynamics that can help interpret our results. The baseline agent persevering on the first target it found is like an attractor capturing the agent’s behavior. Destabilizing the attractor with asymmetric STDP or making it shallower with many small place fields can encourage exploration. Alternatively, injecting noise can help the agent escape the attractor. Each mechanism has drawbacks: destabilized attractors lead to low performance, shallower attractors result in slower convergence and faster forgetting, and injecting noise requires external signals.

In conclusion, our modeling suggests that intrinsic neural mechanisms may not be sufficient to simultaneously ensure behavioral flexibility, rapid learning, and reasonable performance. A second system might be needed to monitor and intervene in the agent’s navigation and learning.

## Acknowledgments

This work was supported by the Deutsche Forschungsgemeinschaft (DFG, German Research Foundation) – project number 316803389 – through SFB 1280, projects A14. We thank Amany Omar, Paul Adler and Raymond Black for help with the figures and the CoBeL-spike software framework.

## Code availability

The CoBeL-spike framework is freely available available at https://github.com/sencheng/CoBeL-spike.

## Footnotes

1 https://github.com/sencheng/CoBeL-spike

